# Tree Based Co-Clustering Identifies Variation in Chromatin Accessibility Across Hematopoietic Cell Types

**DOI:** 10.1101/2021.05.07.443145

**Authors:** Thomas B. George, Nathaniel K. Strawn, Sivan Leviyang

## Abstract

Chromatin accessibility, as measured by ATACseq, varies between hematopoietic cell types in different branches of the hematopoietic differentiation tree, e.g. T cells vs B cells, but methods that relate variation in chromatin accessibility to the placement of a cell type on the differentiation tree are lacking. Using an ATACseq dataset recently published by the ImmGen consortium, we construct associations between chromatin accessibility and hematopoietic cell types using a novel co-clustering approach that accounts for the structure of the hematopoietic, differentiation tree. Under a model in which all loci and cell types within a co-cluster have a shared accessibility state, we show that roughly 80% of cell type associated accessibility variation can be captured through 12 cell type clusters and 20 genomic locus clusters. Using publicly available ChIPseq datasets, we show that our clustering reflects transcription factor binding patterns with implications for regulation across cell types. Our results provide a framework for analysis of chromatin state variation across cell types related by a tree or network.

## 1 Introduction

The development of the ATACseq technique over the past decade has spurred a broad investigation of chromatin accessibility across cell types [4, 18]. In particular, chromatin accessibility has been intensively studied using ATACseq across many hematopoietic cells types [8, 20, 32, 21, 5, 43, 39, 42]. Hematopoiesis, which involves the differentiation of a single hematopoietic stem cell into the different blood cell types, is well characterized and the lineages through which the differentiation occurs can be described by a differentiation tree [33]. Chromatin accessibility has been shown to vary across different hematopoietic cell types, and these differences have been shown to be essential to cell differentiation and cell function, e.g. [13, 38, 36, 16, 21, 39].

While many studies have shown differences of accessibility across hematopoietic cell types, we lack a quantitative description of how variation in accessibility across the hematopoietic cell types reflects the form of the differentiation tree. In this context, many questions remain unanswered. Are most accessible genomic loci accessible only in a specific, cell type or are a significant number of loci accessible jointly across a particular collections of cell types (e.g. all T cells)? If many loci are accessible across a particular collection of cell types, do those cell types form connected components of the differentiation tree, or are they dispersed? More generally, how do we quantitatively find and describe associations between chromatin accessibility and the differentiation tree? And finally, if such associations exist, how do they shape cellular regulation? Answering these questions would provide a context in which to analyze the chromatin accessibility of particular hematopoietic cell types and to understand differences in cellular regulation across the hematopoietic cell types.

Here, we address these questions using a recently completed ImmGen ([35]) consortium dataset published by Yoshida et al. [43]. Yoshida et al. characterized accessibility through bulk ATACseq across 90 murine, immune cell types. The availability of bulk ATACseq across such a large number of cell types using consistent protocols provides a novel opportunity to investigate accessibility patterns in hematopoietic cell types. Typically, in ATACseq studies across multiple cell types, the ATACseq workflow ends with the formation of an accessibility matrix *M*, with the rows of *M* corresponding to genomic loci and the columns corresponding to cell types. *M* can be binary, reflecting a call of accessible or not-accessible for a particular genomic locus in a particular cell type or can take a range of values, for example if the height of the ATACseq peak is used to quantify accessibility. Describing chromatin accessibility across cell types can then be framed as describing the structure of *M*. In our context, we are interested in understanding how the structure of *M*, which is built from the Yoshida et al. ATACseq dataset, reflects the hematopoietic differentiation tree.

In a general setting, the most common way to describe the structure of a matrix, *M*, is to construct another matrix, 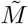, with some simple form that is a good approximation of *M*. Standard approaches, such as the svd, are difficult to interpret, and have not been commonly used in the context of genomics data. Starting with gene expression datasets in the early 2000s [11, 26, 29] and extending to current ATACseq datasets, clustering has been the most common approach to describing *M*.

In the case of chromatin accessibility datasets, a common clustering analysis involves clustering of the columns (i.e. cell types), typically by dimension reduction followed by k-means or by hierarchical clustering, which identifies cell types with similar chromatin accessibility patterns across the genome, e.g. [7, 31, 9]. Column clustering has the advantage of decomposing the cell types of the differentiation tree into distinct clusters that can then be analyzed. However, cell type clustering provides little information about the overall structure of *M* which typically has many more rows than columns. Row (i.e. locus) based clustering, which identifies loci with similar cell type accessibility patterns, is also common, e.g. [38, 20, 32, 21, 43]. Lara-Astoria et al., [20], used k-means to row cluster a dataset involving 16 hematopoietic cell types. They noted that loci in different row clusters were accessible across different cell types (see their Figure 3). For example, in one of their locus clusters, the loci were accessible in stem cells, but not in other cell types. Yoshida et al. used t-SNE to project rows (i.e. loci) onto 2-d and then identified loci that cluster in the 2-d space and are either accessible across all cell types or are accessible within a single cell type. Both these examples reflect an association between *M* and the differentiation tree, but restriction to row clustering limits investigation of the association.

Biclustering, which involves specifying a joint cluster of rows (i.e. loci) and columns (i.e. cell types), received significant attention during the 2000s in the context of gene expression data [6, 22, 34]. Biclustering is particularly effective at finding substructure within *M* and multiple biclusters can be found and used to construct an approximating matrix 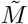, but the resulting approximating matrix 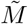 can be difficult to interpret and its computation is often unstable [27].

Here, we take a middle ground between row or column based clustering and biclustering by considering the structure of *M* through co-clustering. By co-clustering, we mean selecting locus clusters and column clusters whose pairings provide a grid-like structure to the approximating matrix 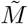. In some sense, our work is an extension of the row clustering results of Lara-Astasio et al. (discussed above) which suggested that the structure of *M* can be well approximated by co-clustering. Our approach is to first row cluster, using the well-known Louvain algorithm, [3], which allows for scaling to large number of loci, a typical situation in chromatin studies. We then column cluster. But importantly, we develop a novel clustering algorithm that restricts column clustering to clusters that are composed of coherent components of the hematopoietic differentiation tree. This co-clustering provides a simple and biologically meaningful structure to 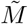 in which a particular column cluster is a coherent hematopoietic phenotype and the overall structure of *M* can be viewed through the accessibility of these hematopoietic phenotypes across multiple locus clusters. Further, the construction allows us to characterize the variance in *M* that is associated with the differentiation tree.

Previous authors have considered clustering loci in the context of a cell type network such as the hematopoietic differentiation tree, but typically with the goal of annotating loci [2, 44, 37]. For example, treeHMM [2] infers a hidden state at each genomic locus for each of the cell types through a hidden Markov model, with the hidden state serving as an annotation of the locus. From our perspective, these methods serve to construct a matrix *M* - which is the starting point of our analysis - with the value of *M* being the hidden state of the HMM across cell types and loci. In contrast to our setting, in which we just have ATACseq data, these methods allow for multiple assays for each cell type - for example ChIPseqs of different histone modifications in which case constructing *M* is complex.

We show that roughly 1*/*2 of accessible loci in the Yoshida et al. dataset are accessible in only one or two cell types in the differentiation tree. Putting aside these cell type specific loci, we show that the other loci fall into roughly 20 locus clusters. Each of these locus clusters can be characterized by cell types in which the accessibility is relatively high and cell types that are relatively low, and the cell types with high accessibility compose a coherent component of the differentiation tree. We show that with 12 cell type clusters (i.e. column clusters) that decompose the differentiation tree, we can capture roughly 80% of the cell type specific variation in *M*. We also investigate transcription factors (TFs) in the context of this co-clustering, showing that the co-clustered structure of *M* is reflected in the motif and binding patterns of TF across loci and cell types.

## 2 Results

We downloaded the ATACseq data of Yoshida et al. from the NCBI GEO database [1]. The Yoshida et al. dataset includes ATACseq libraries across 90 cell types, but we restricted our attention to 78 of the cell types, leaving out some outlier cell types. The differentiation tree we assume is shown in Figure 1 and is the same as that of Yoshida et al. (compare to their Figure 1A) but without the outlier cell types. We applied the ENCODE ATACseq pipeline [10] to the Yoshida et al. ATACseq samples for each of the cell types, resulting in a collection of peaks representing accessible loci for each cell type. Then, following the approach of previous authors, e.g. [20, 8, 43], we constructed a master list of loci, composed of non-overlapping 250 base pair windows that intersected with every ATACseq peak across all cell types. We then formed the chromatin accessibility matrix *M* with the rows corresponding to each of the loci in the master list and the columns corresponding to the 78 cell types. An entry of *M* was 1 if the cell type had a peak that intersected with the corresponding 250 base pair locus, otherwise the entry was 0.

**Figure 1:**
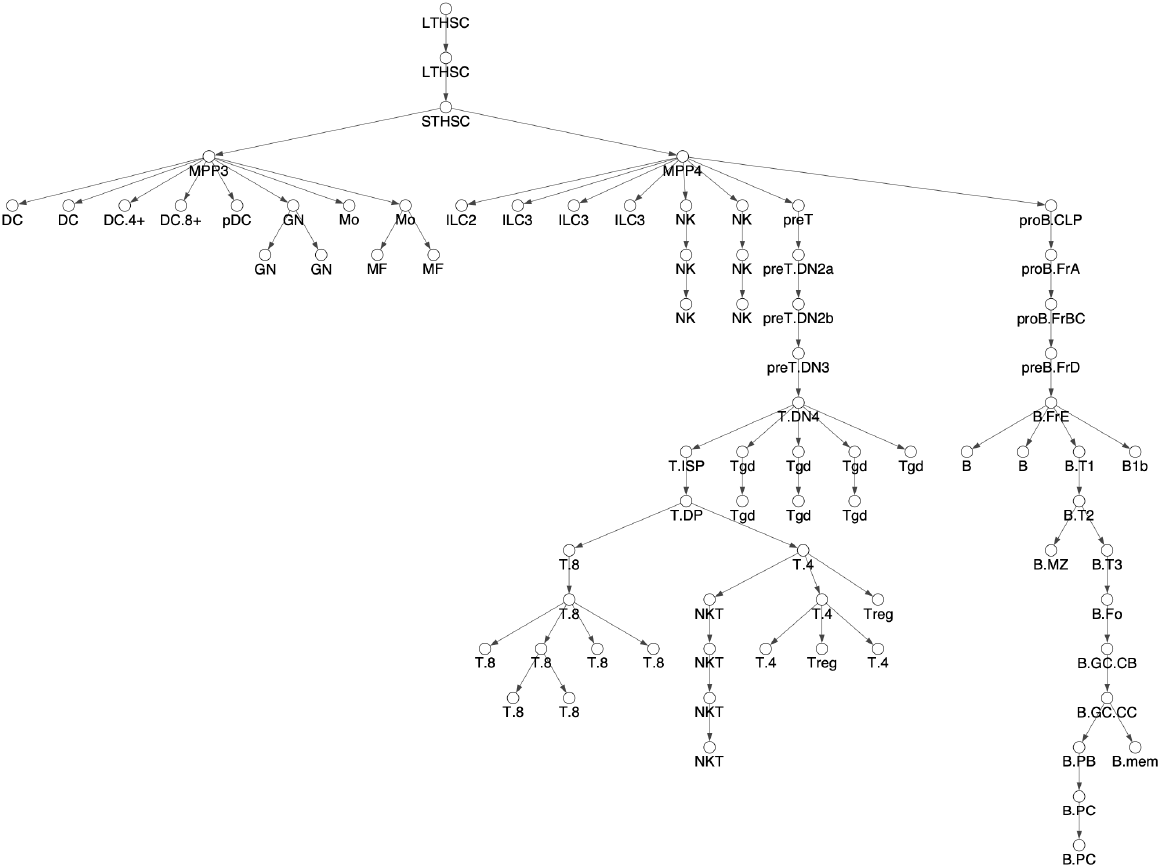
Differentiation Tree. We consider 78 hematopoietic cell types and assume the shown tree structure. The tree structure is essentially the same as Fig 1A in Yoshida et al. Cell type names have been shortened for readability.

An important issue in constructing *M* is the sensitivity, specificity trade-off in calling locus accessibility. We modulated this tradeoff by choosing different IDR values [23] in the ENCODE pipeline. The IDR is similar to an FDR and is used to determine reproducible peak calls. We considered IDR values of 0.01, 0.05, 0.10 and 0.15 which led to roughly 159, 246, 331, and 424 thousand loci which were accessible, respectively, in one or more of our 78 cell types. As comparison, Yoshida et. al. considered roughly 512 thousand loci over their 90 cell types. We defined a cell specific locus as a locus which was accessible in 2 or less cell types; we found that 38%, 43%, 51%, and 57% of the loci for the respective IDR values were cell specific. The increasing level of cell specific loci with increasing IDR may reflect increasing noise. Alternatively, cell specific accessible loci may have lower levels of accessibility, leading to their being called only at larger IDR values. Regardless of the specific IDR, the number of cell specific and non-cell specific loci both constituted a substantial fraction of the total accessible loci. Below we present results for an IDR of 0.01, taking a conservative approach to calling accessibility. Our results were essentially unchanged using other IDR, see Materials and Methods for further details.

### 2.1 Row Clustering

We clustered the rows (i.e. loci) of the accessibility matrix *M* using the Louvain algorithm [3]. Importantly, we only clustered the rows corresponding to non-cell specific loci, of which there were 98, 848 under 0.01 IDR. The Louvain algorithm takes a graph (i.e nodes and edges) as input and clusters the nodes to maximize a measure of community structure. In our setting, the rows (i.e. loci) of *M* form the nodes. To form edges, we placed an edge between two nodes if the corresponding rows had a statistically significant number of columns with equal entries (i.e.the two loci shared accessibility states across a statistically significant number of cell types). We clustered nodes with edges placed at an FDR of 0.0001, 0.001, 0.01, 0.05, and 0.10, respectively. As shown in Table 1, the fraction of nodes connected by an edge to some other node fell as edge FDR was lowered, reflecting the existence of loci with cell type accessibility patterns that did not closely match any other loci. As the table further shows, the Louvain algorithm formed between 16 to 21 clusters with 30 or more nodes across all FDR, except in the case of an FDR of 0.0001. The results for an edge FDR of 0.0001 suggest an overly conservative approach in placing edges, leading to too many clusters and many isolated nodes. Below, we present results for the edge FDR of 0.001, choosing a relatively conservative value as we did for the IDR. Our results are essentially unchanged using the larger edge FDR values, see Materials and Methods for details.

**Table 1:**
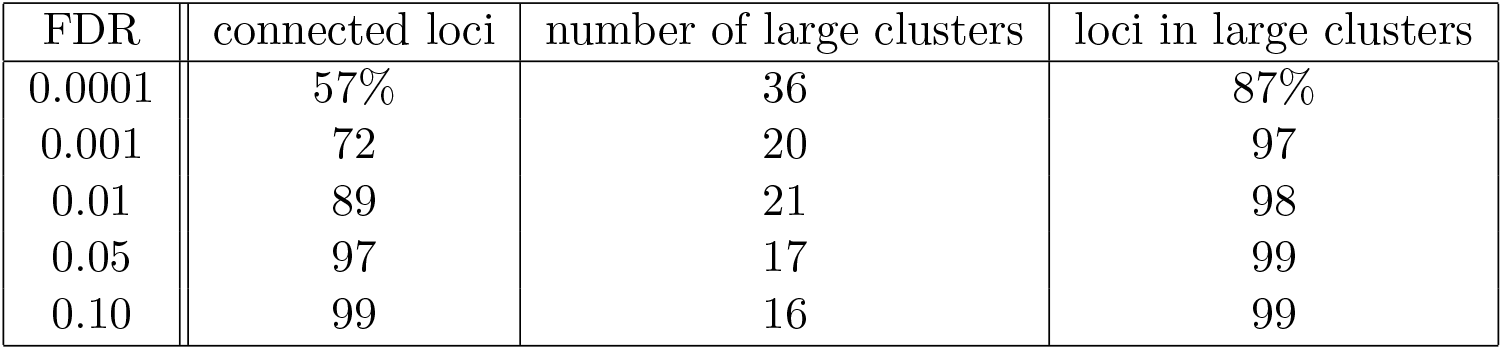
The Effect of Edge FDR on Locus Clustering. To apply the Louvain clustering algorithm, we constructed a graph in which loci were represented by nodes and edges between nodes represented loci with similar accessibility patterns. We placed edges between two nodes at different FDR. Shown are the percent of loci that were connected to another locus (connected loci), the number of clusters with more than 30 loci (number of large clusters), and the fraction of loci that fell within a large cluster (loci in large clusters).

At an edge FDR of 0.001, the Louvain algorithm produced 100s of clusters, but the top 20 clusters included 97% of the nodes in the graph with the remaining clusters all containing less than 30 nodes and most containing only 2 nodes. The smaller clusters could reflect noise in calling loci and edges or they could reflect loci with uncommon patterns of accessibility. Table 2 shows the number of loci in each of the largest 20 locus clusters. The first 13 clusters have greater than 2000 loci and the rest of the clusters have 100s of loci, except for cluster 19 which has 45 loci. Figure 2 shows the fraction of loci in each row cluster that are accessible within each of the 78 cell types.

**Table 2:**
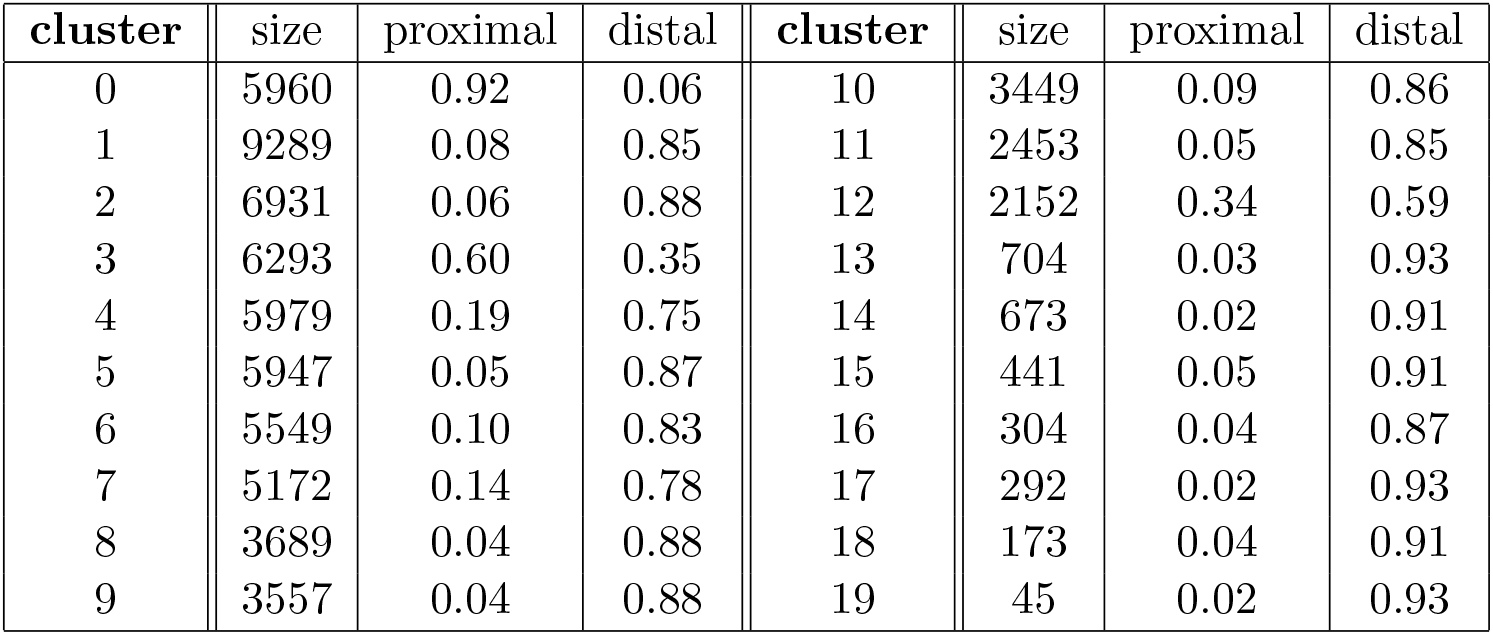
Row Clusters. Twenty row clusters generated by the Louvain clustering contained 95% of the non-cell specific loci. The twenty row clusters had varying number of loci (size) and a different fraction of loci within 500 base pair (proximal) and more than 3000 base pair (distal) from of a transcription start site (TSS). Row cluster 0 was unique in having loci largely proximal to TSS, possibly acting within promoters. All other clusters were largely composed of loci distal to TSS, possibly acting within enhancers.

**Figure 2:**
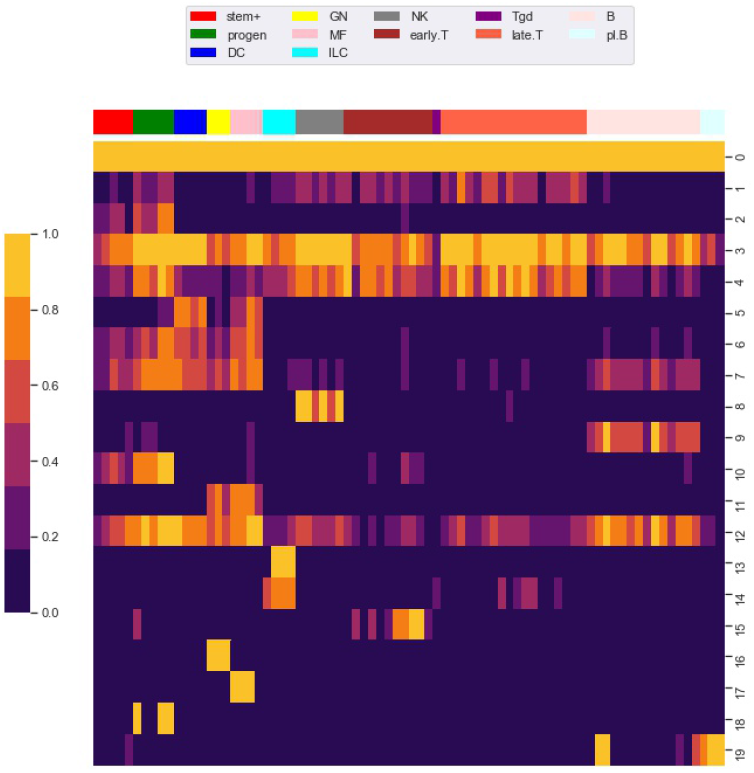
Row Clustering using the Louvain Algorithm. We used the Louvain algorithm to cluster 98, 848 rows (i.e. loci) of the *M* matrix, giving 20 main row (i.e. locus) clusters. Each row in the heat map corresponds to a row cluster and each column to a cell type. The number of loci in each row cluster is given in Table 2. Heatmap colors show the fraction of loci within a row cluster that were accessible for the particular cell type. Column groupings by cell type are based on our column clustering for *k* = 12 discussed below.

Notably, cluster 0 contains loci that are accessible in almost all cell types. As shown in Table 2, most of the loci in this cluster are within 500 base pair of a transcription start site (TSS), possibly acting within promoters. Most of the loci in the other locus clusters, accessible in only a subset of the cell types, were greater than 3000 base pair from a TSS, possibly acting within enhancers. Yoshida et al. noted a similar pattern, with loci close to TSS (what they term TSS OCR) accessible across most cell types and loci far from TSS (what they term DE OCR) accessible in only certain cell types.

The clustering reveals clear associations between cell phenotype and accessibility. As an example, Figure 3 shows the fraction of loci in row clusters 5, 7 and 9 called as accessible for each cell type. The figure gives the same data as rows 5, 7 and 9 of the heatmap in Figure 2, but in the context of the differentiation tree. In row cluster 5, roughly 50% − 80% of loci are accessible in macrophages and DCs, while in other cell types accessibility of these loci is less than 1%. In row cluster 7, between 40% − 80% of loci are accessible in macrophages, dendritic cells, and most B cells, while in other cell types less than 10% of loci are accessible. In row cluster 9, all B cells except pro-B cells and plasma B cells have between 60% − 95% of loci as accessible, while in all other cell types less than 10% are accessible. These three row clusters reflect a decomposition of accessible loci in macrophages, DC and B cells into loci that are accessible only in macrophages and DC, only in B cells and jointly. Some of the smaller row clusters, which reflect a more specific cell phenotype, have a more definitive separation of accessibility across cell type. For example, all 704 loci in cluster 14 are accessible in ILC3 cell types and inaccessible in all other cell types. The less definitive separation in cell type accessibility we see in clusters 5, 7 and 9 may reflect experimental noise in the ATACseq workflow, but particular loci within a cell type may also vary in their accessibility over time due to unstable positioning of nucleosomes or due to a variation of cell state [41, 24].

**Figure 3:**
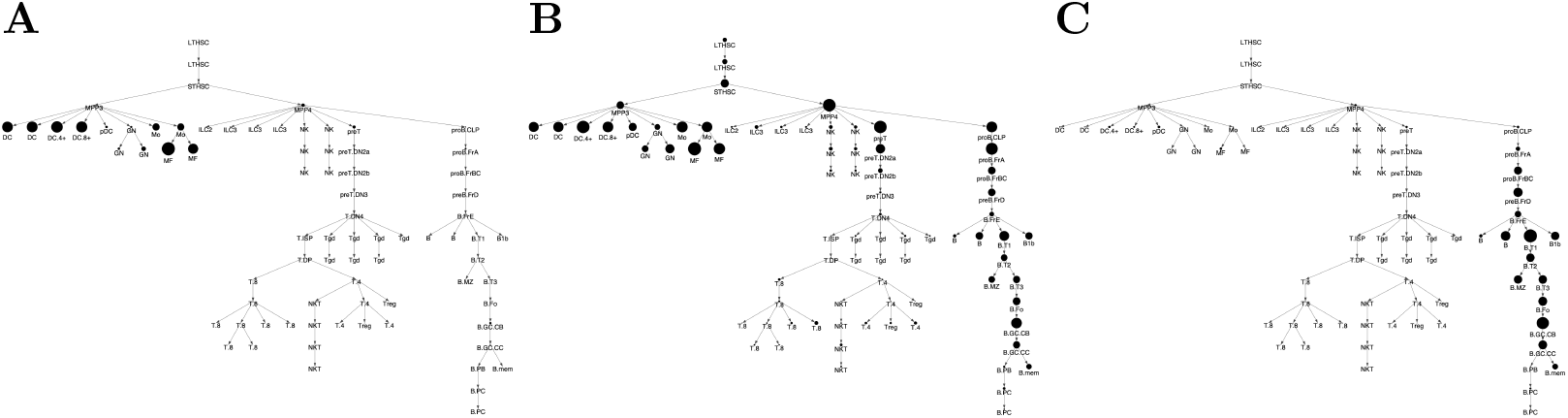
Locus Accessibility Across Locus Clusters Reveals Cell Type Specific Patterns. Shown is the fraction of loci that are accessible within a locus (i.e. row) cluster across each of the cell types in the differentiation tree. Trees correspond to locus cluster (A) 5, (B) 7, and (C) 9. Within each tree, the size of a vertex provides the fraction of loci accessible for the corresponding cell type. Figure 2 provides the same data, but in a heatmap format.

### 2.2 Cell Type Clustering

With the row clustering fixed, we next applied a column (i.e. cell type) clustering. For column clustering, we chose the number of column clusters as a particular value, *k*, and produced a column clustering for each *k* = 2, 3, *…*, 12. Since one of our main motivations was to quantify the degree to which accessibility associates with the differentiation tree, we restricted column clusters to reflect the structure of the tree.

Given a graph (i.e. nodes and edges), a collection of nodes is said to be connected if there is a path along the tree from every node to every other node. Importantly, restricting cell type clusters to connected components did not give good results. As an example, consider accessibility in locus (i.e. row) cluster 5, as shown in Figure 3. Only 5% of loci in the cluster are accessible in the MPP3 progenitor cell type, but 80% the loci are accessible in macrophage and DC, which are children of the MMP3 cell type. Further, for pDC (plasmacytoid DC) and GN (neutrophil), which are also children of MMP3, the loci are relatively inaccessible (*<* 30%). To account for this dynamic, we say a cell types cluster *respects* the differentiation tree if the nodes in the cluster can be divided into a collection of connected components, and that these connected components each descend from a particular parent node which need not be part of the cluster. For a given *k*, our clustering algorithm selected the cell type clustering that respected the tree and led to the best co-clustering approximation of *M*, see Methods for details.

Figure 4 shows the column clustering produced for *k* = 3, 8, 12 on the differentiation tree. When *k* = 3, our clustering splits the differentiation tree into a stem/progenitor cell and myeloid cluster, a B cell cluster, and a T cell cluster. This clustering shows that the general division of immune cell types into myeloid, B, and T phenotypes is reflected in accessibility differences. Further, stem cell and progenitor cell types are most similar to myeloid cell types in their accessibility. Increasing to *k* = 8, splits stem cells and progenitor cells into separate clusters, puts the NK and ILC cell types into separate clusters, introduces a cluster containing plasma and memory B cells, and splits the myeloid compartment. Interestingly, neutrophils are grouped with stem cells, while DC, monocytes, and macrophages are split into their own compartment. By *k* = 12, the T cell compartment is split into a CD8 and CD4 cluster and an early T cell cluster.

**Figure 4:**
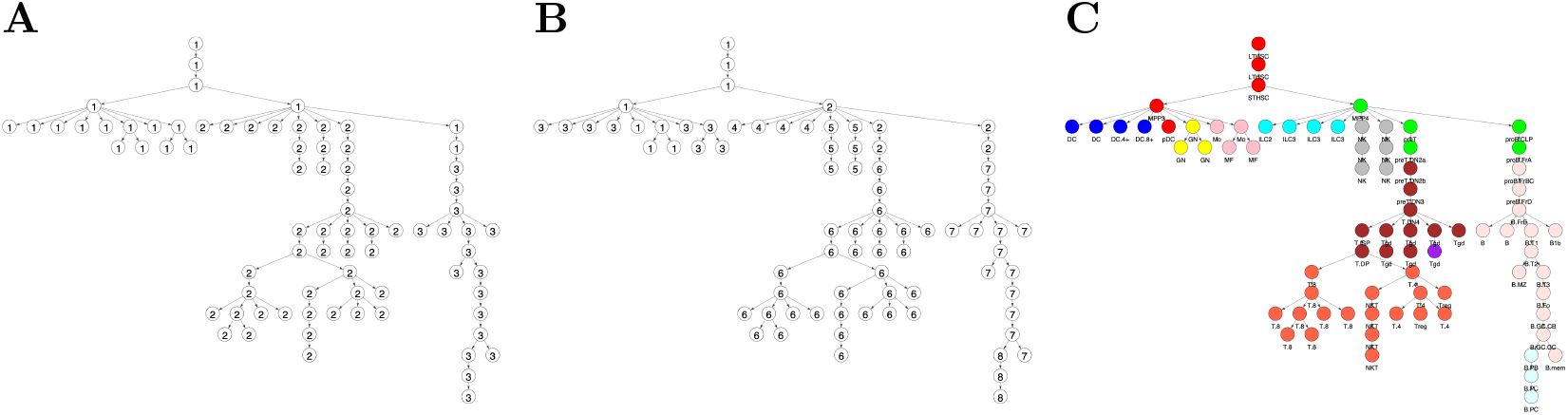
Cell Type Clusterings that Respect the Hematopoietic Differentiation Tree. Shown are the column (i.e. cell type) clusters for (A) *k* = 3 (B) *k* = 8; and (C) *k* = 12, which is the cell type clustering we use throughout the text. In panel C, clusters are specified by color and this color scheme is used throughout the text. In panels A and B, clusters are specified by number.

### 2.3 Co-clustered Approximation of the Accessibility Matrix

For a particular *k* value, the combination of row and column clusters divided *M* into a grid of 20 × *k* co-clusters. While *M* is a binary matrix, we built the co-clustered approximation of *M*, 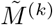, by setting all the entries in a co-cluster to the mean value of the co-cluster entries in *M*. In constructing the matrix 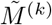, we allowed each co-cluster to take on a different value. As a more restrictive model, we assumed that a cell type within a particular row (i.e. locus) cluster can be either in an relatively inaccessible (i) or accessible (a) state, rather than in a continuum of accessibility states. Biologically, this more restrictive model supports a single regulatory mechanism shaping the accessibility structure of the loci in each locus cluster. To examine the impact of this model, we built a matrix 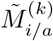 which had the same co-clusters as 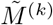, but with co-clusters sharing the same locus cluster restricted to have one of two values, representing either an inaccessible or accessible state. Figure 5 visualizes these two matrices for *k* = 12.

**Figure 5:**
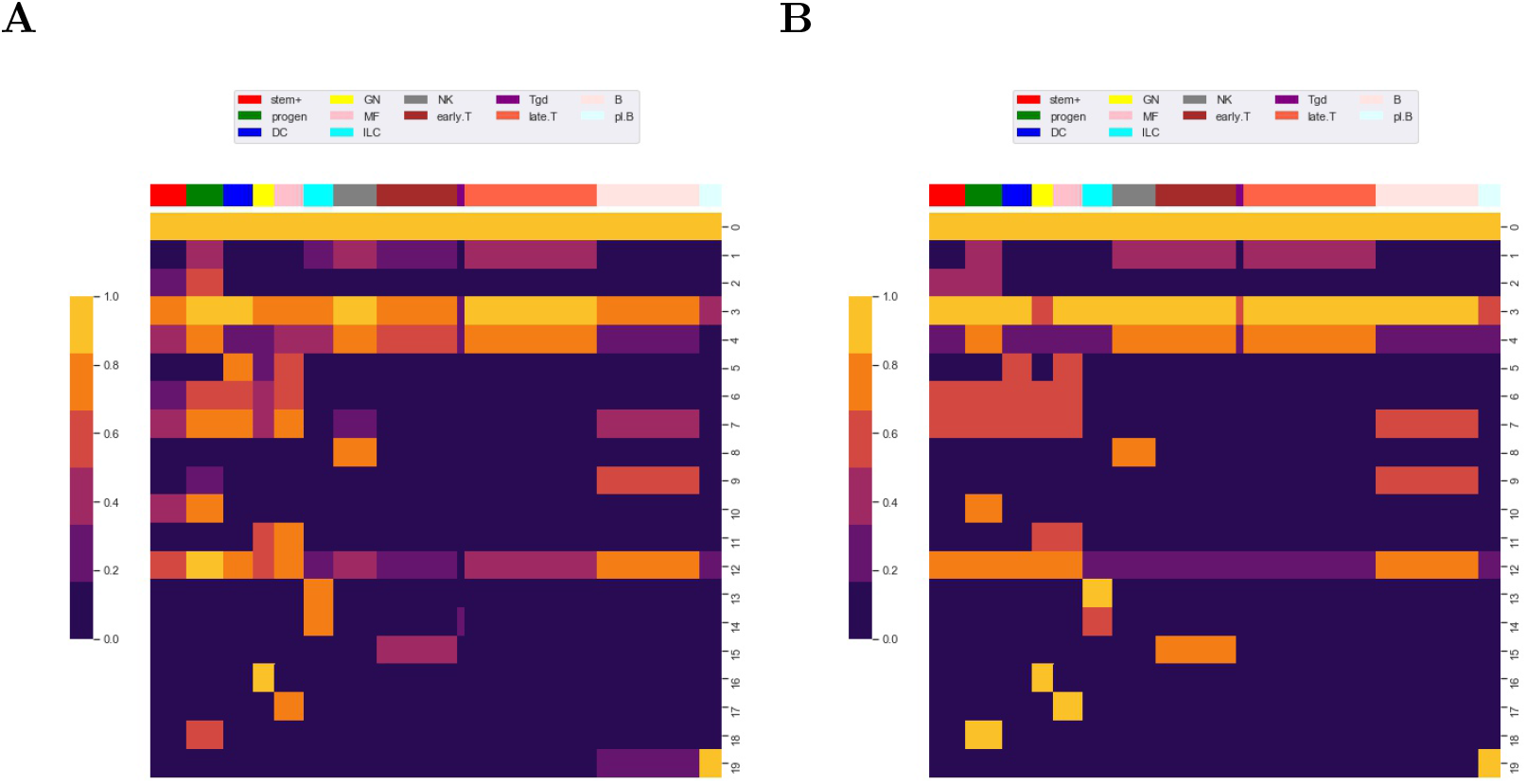
Co-clustered Approximation of the Accessibility Matrix *M*. Heatmaps of (A) 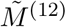 and (B) 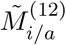 The co-clustered matrices 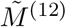 and 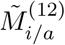 approximate *M*. Compare the heatmaps of this figure to the heatmap of *M* given in Figure 2.

We applied an ANOVA analysis to calculate the variation of *M* captured by the co-clustered matrices 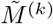 and 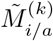. We calculated two R-squared values,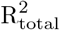 and 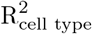, for the fraction of the total variation in *M* captured by the co-clustering and the fraction of cell type (i.e. column) associated variation in *M* captured by the cell type clustering, respectively; see Methods for details.If each cell type was in a separate cluster, then 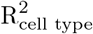 would equal 1 and if all cell types were in a single cluster then 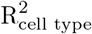 would equal 0. Since we constructed our column clusters to respect the differentiation tree, we used 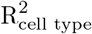 as a measure of the association between accessibility and the structure of the differentiation tree. Figure 6 shows the R-squared values for different values of *k*. Also included in the figure are co-clusterings in which cell types were clustered using hierarchical and kmeans clustering, respectively. We used the R-squared values for these two general clustering methods, which are not constrained by the differentiation tree, as a baseline against which to compare the R-squared of our clustering approach, which is constrained by the tree.

**Figure 6:**
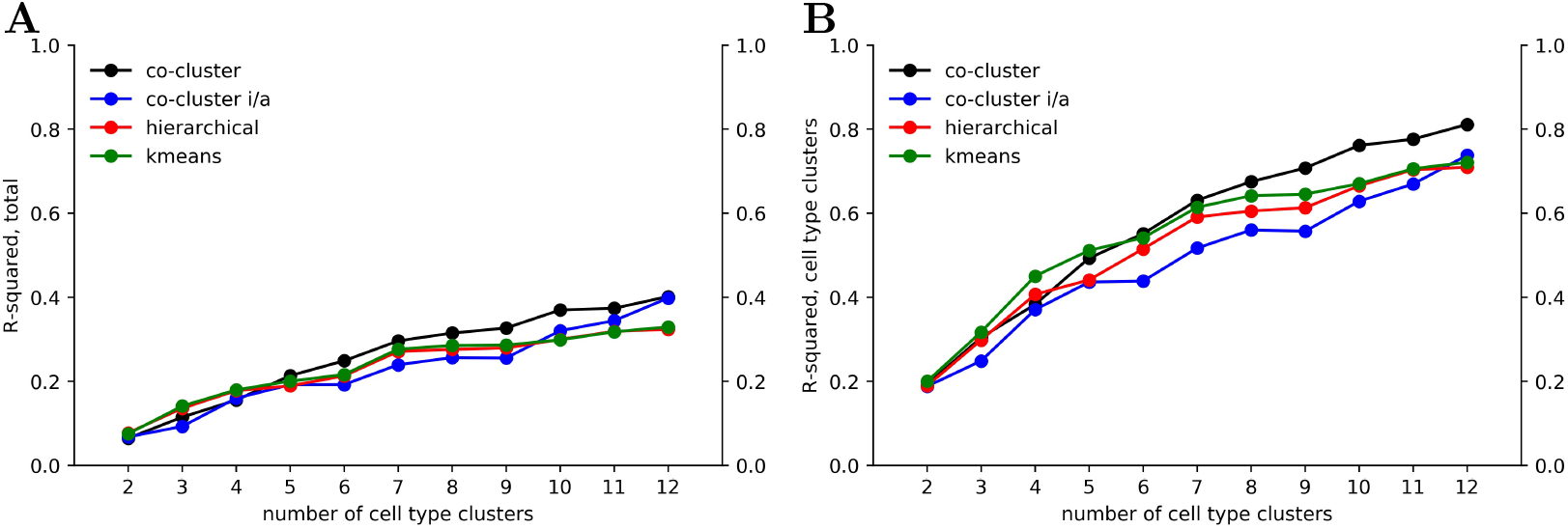
Decomposition of Accessibility Variation. Using an ANOVA analysis, we calculated an R-squared value giving the fraction of (A) total variation and (B) cell type associated variation in *M* captured by our co-clustered approximation matrices 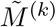 and 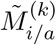 as well as co-clustering in which columns were clustered using hierarchical and k-means methods without taking the differentiation tree into account.

As seen in Figure 6A, the fraction of variation captured by the co-clustering varied between 0.20 to 0.40 as the number of cell type clusters *k* rose from 2 to 12. In contrast, as shown in Figure 6B, the fraction of cell type associated variation captured by the cell type clusters rose from 0.20 to 0.80, meaning that our cell type clustering captured most of the association between accessibility and cell type, at least for the higher *k* values. The large fraction of the total variation in accessibility not captured by our co-clustering (roughly 0.6 for *k* = 12) is associated with the row clustering. As discussed above and reflected in Figure 2, within a row (i.e. locus) cluster, cell types tended to have either a high or low fraction of loci in an accessible state, but the high and low fraction were often intermediate, e.g. 0.50 and 0.10, instead of extreme, e.g. 1.0 and 0. This within cell type variation could reflect noise in the ATACseq workflow, stochasticity in the accessibility state of the loci, or row clusters that are too broad.

The 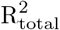 and 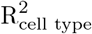 values based on the 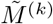 co-clusterings were similar to values based on 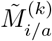 and hierarchical and kmeans column clusterings. The similarity of the R-squared values between 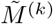 and 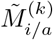 provides support for viewing accessibility within a particular locus (i.e. row) cluster as falling into one of two states for all the cell types. The clusters inferred by the hierarchical and kmeans algorithms included clusters with cell types dispersed through the tree, but this increased degree of freedom did not associate with a better co-clustering approximation. The similarity of the R-squared values based on hierarchical and kmeans clusterings to our co-clusterings suggests that there is not a significant component of cell type variation that does not respect the differentiation tree.

### 2.4 TF Motif Enrichment Across Co-Clusters

Using methods introduced by Schep et al. in [30], Yoshida et al. computed an accessibility score measuring the enrichment of a TF motif on accessible loci within a particular cell type relative to the presence of the TF motif on accessible loci across all cell types. From the perspective of a matrix analysis, this is a column (i.e. cell type) based approach because enrichment is assessed over all accessible loci for a single cell type. To adapt the method of Schep et al. to co-clusters, we defined 20 co-clusters formed by the combination of each locus cluster and the cell types in the accessible state of 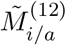 for that locus cluster, see Figure 7. We refer to these co-clusters as accessibility co-clusters. There is one accessibility co-cluster for each locus cluster. For example, accessibility co-cluster 9 is formed by the loci in locus cluster 9 and the cell types in our B cell, cell type cluster while accessibility co-cluster 5 is formed by the loci in locus cluster 5 and the DC and macrophage cell type clusters. For each accessibility co-cluster, we adapted the method of Schep et al. by considering enrichment of a TF motif over accessible loci in the accessibility co-cluster against a background of accessible loci over all other loci and cell types, see Materials and Methods for computational details.

**Figure 7:**
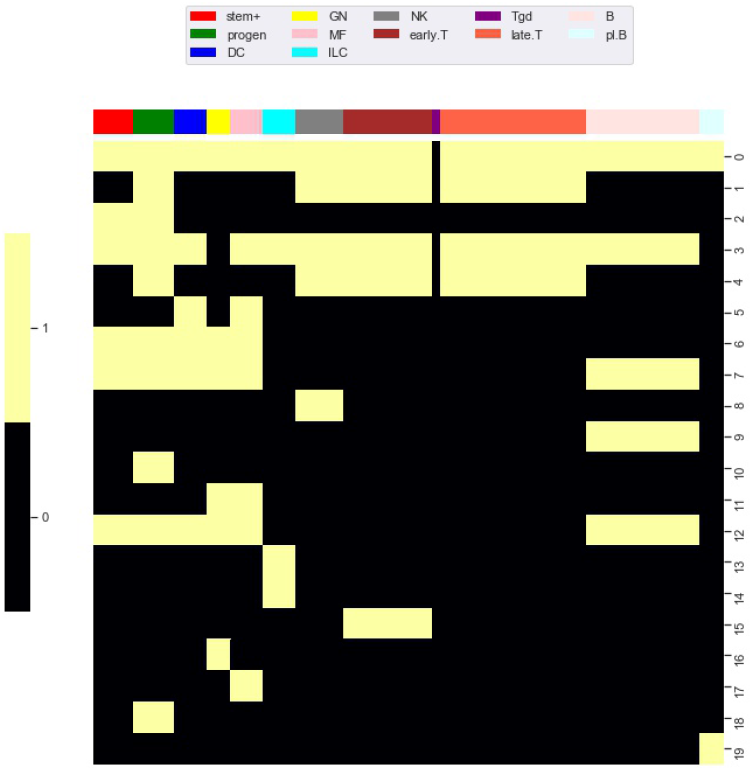
Accessibility Co-Clusters. To investigate enrichment of TF motifs, we defined a collection of accessibility co-clusters formed through the combination of locus clusters and all cell types in the accessible state of 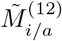. In the heatmap, accessibility co-clusters are the yellow blocks within each of the 20 rows, compare to 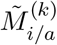 in Figure 4B. As an example, accessibility co-cluster 5 is formed by the loci in locus cluster 5 and the DC and macrophage cell type clusters

Importantly, since loci near TSS tended to be accessible across all cell types, we restricted our analysis to loci greater than 3000 base pair from a TSS. This had the advantage of restricting our analysis to regulatory features specific to putative enhancers, which are likely different than regulatory features specific to promoters. For TF motifs, we used the 76 TF motifs identified by Yoshida et al. as significantly associated with accessibility (see their Table S5). Of these TF motifs, we found that 43 were statistically enriched in at least one of the 20 accessibility co-clusters (FDR 0.05).

Figure 8 shows enrichment across the 43 significant motifs and the 20 accessibility co-clusters. Of the 43 TF motifs that we found to be enriched in at least one accessibility co-cluster, the 7 motifs BCL11A, BCL11B, NFE2, NFKB1, RUNX1, RUNX2, RUNX3 were enriched in more than 3 accessibility co-clusters. The TF BCL11A is an instructive example. Yoshida et al. found BCL11A motifs enriched in accessible loci in B cells and myeloid cell types (see their Fig 5F). Similarly, we find BCL11A to be enriched in 4 accessibility co-clusters: the co-cluster formed by locus cluster 5 and myeloid cell types; the co-cluster formed by locus cluster 7 and stem cells, myeloid cells and B cells; the co-cluster formed by locus cluster 9 and B cells; and the co-cluster formed by locus cluster 12 and myeloid cells and B cells. Our enrichment analysis extends the results of Yoshida et al. by decomposing the cell type enrichment of BCL11A. For example, the enrichment of BCL11A in accessible B-cell loci reflects enrichment in loci that are jointly accessible across myeloid and B cells, but also in loci that are accessible only within B cells or myeloid cells, respectively.

**Figure 8:**
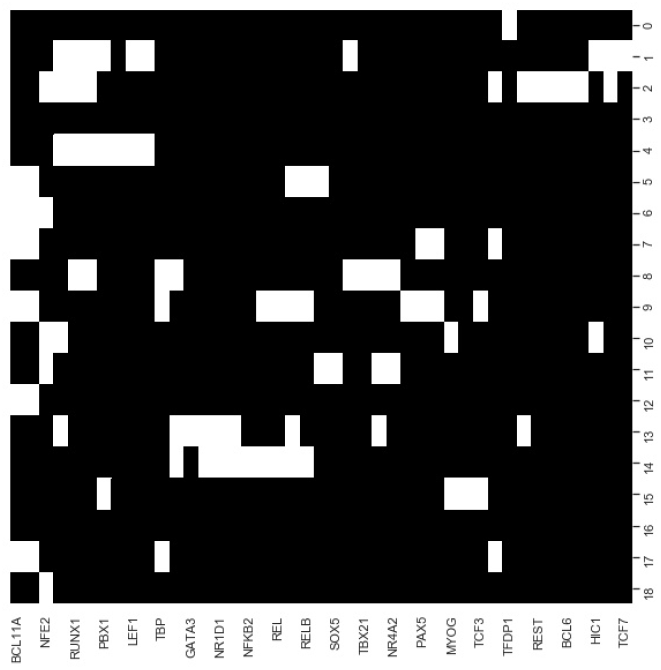
TF Motif Enrichment Associates with Particular Accessibility Co-clusters. We evaluated enrichment of each TF motif (columns) within each accessibility co-cluster (rows). At an FDR of 0.05, we found 43 TF motif, co-cluster pairings that were enriched (white coloring). As a particular example, the BCL11A motif (far left column) was enriched in 6 accessibility co-clusters, in particular cocluster 9. Accessibility co-cluster 9 is formed by loci in row cluster 9 and B cell types. The co-cluster is visualized in Figure 7 by the yellow block within row 9.

There were 13 motifs enriched for a single accessibility co-cluster with the most significant enrichment occurring for TCF12, TBX21, NFKB2, EBF1, and GATA3. EBF1 and GATA3 are instructive examples. Yoshida et al. also generated RNAseq datasets for each of their cell types. Based on these RNAseq datasets, EBF1 is expressed solely in B cells while GATA3 is expressed in ILC, NK, and T cells. Reflecting these expression patterns, EBF1 is known as a master regulator of B cell differentiation [25] and GATA3 is a regulator of T cell differentiation [14]. We found the EBF1 motif enriched in the co-cluster formed from locus cluster 9 and B cell types, matching the known regulation role of EBF1. In contrast, GATA3 was enriched in the co-cluster formed from locus cluster 13 and ILC3 cell types. This result matched at least part of the expression pattern of GATA3, but did not reflect the regulatory role of GATA3 in T cell differentiation. For GATA3, our enrichment shows that ILC3 cells have accessible loci that are more enriched for the GATA3 motif than T cells, but the regulatory significance of this result is unclear.

The remaining 23 TF were enriched in 2 or 3 co-clusters. The transcription factors PAX5 and EOMES are instructive examples. PAX5 is a master regulator of B cell differentiation [15]. We find PAX5 enriched in the co-clusters formed by locus cluster 7 and myeloid and B cells and by locus cluster 9 and B cells. Yoshida et al.’s RNAseq data show that PAX5 is expressed solely in B cells, so PAX5 motifs in locus cluster 5 are not bound by PAX5 in myeloid cell types, but these loci are accessible, suggesting an association with other TFs. EOMES regulates effector NK and T cells [12] and, in line with this regulation, Yoshida et al.’s RNAseq data shows EOMES expressed in NK cells and CD8 T cells. We find EOMES enriched in two accessibility co-clusters, one formed by locus cluster 1 and T cells and one formed locus cluster 8 and NK cells. EOMES motifs are enriched in two locus clusters with loci that accessible in disjoint cell clusters, T and NK cell types, respectively. In contrast, PAX5 motifs were enriched in co-clusters that spanned multiple cell clusters, e.g. B cells and myeloid cells, over which the loci were jointly accessible.

### 2.5 Association of TF ChIPseq Peaks and Co-Clusters

To validate and explore the functional consequences of our TF motif analysis, we collected publicly available ChIPseq datasets from the GEO database for PAX5 and EOMES. We used PAX5 ChIPseq data from Revilla-I-Doming et al., [28], that sampled pro-B cells and mature B cells, and we used EOMES ChIPseq data from Wagner et al. [40] and Istaces et al. [17] that sampled NK cells and CD8 thymocytes, respectively. Briefly, we downloaded fastq files from GEO and applied a standard peak calling workflow to call peaks. We then identified intersections between our master collection of 159 thousand accessible loci and the TF peaks, see Materials and Methods for accessions and workflow details.

Figure 9A shows a scaled count of the number of loci that contained a PAX5 peak across different locus clusters for the mature-B and pro-B cell types. Since locus clusters differ in size, the raw count is not directly informative. Instead, we scaled the raw count by the expected count under the null of equal distribution of peaks across loci. A scaled count above one represents an enrichment of peaks in the locus cluster. Only locus clusters with a scaled count greater than 1 for either the mature B or pro-B ChIPseq datasets are shown. Our TF motif analysis showed enriched motifs in accessibility co-clusters 7 and 9 and, correspondingly, for both the pro-B and mature-B cell ChIPseqs, locus clusters 7 and 9 had enriched counts.

**Figure 9:**
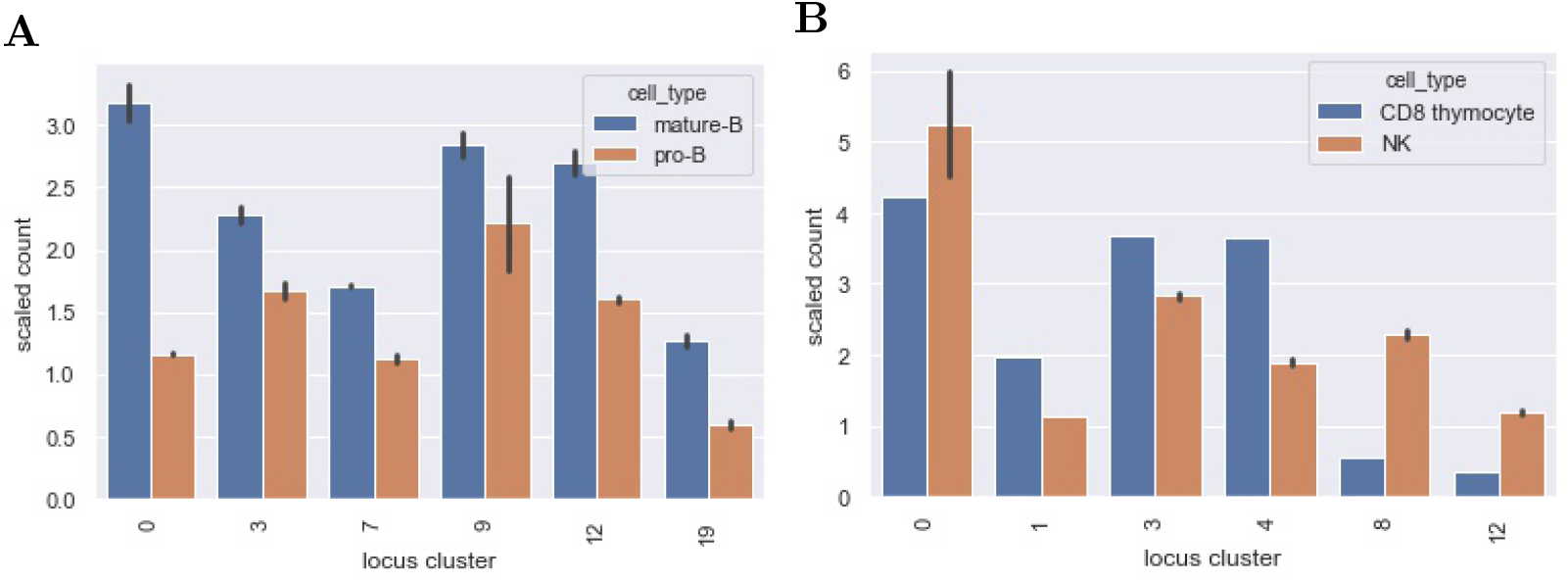
Distribution of PAX5 and EOMES ChIPseq Peaks Across Locus Clusters. The bar graphs show a scaled count (y-axis) of the number of loci in a locus cluster (x-axis) that contained a ChIPseq peak for (A) PAX5 and (B) EOMES ChIPseq datasets. Scaled counts took into account the different sizes of our 20 locus clusters. A scaled count above 1 represents an enrichment of peaks above an expectation under a uniform null. The locus clusters enriched for PAX5 ChIPseq peaks had a statistically significant association with accessibility in pro-B and mature B cell types and we found an analogous association for EOMES peaks, see text for details.

The mature B cell type falls within our B cells, cell type cluster. B cells are present in the accessibility clusters formed by locus clusters 0, 3, 7, 9 and 12 (see the B cell columns of Figure 7). All of these locus clusters had enriched PAX5 peak counts, and only locus cluster 19 had an enriched count without associated accessibility in B cells. Overall, there was a statistically significant association between PAX5 peaks and accessibility (p-value 0.001,, hypergeometric test). The pro-B cell type falls within our progenitor (pro) cell type cluster. Progenitor cells are present in 10 of the accessibility clusters and PAX5 peaks had enriched counts in 4 of these, reflecting a marginally significant association between peaks and accessibility (p-value 0.08). Interestingly, in locus cluster 19, PAX5 peak counts were enriched in the mature B cell ChIPseq but not the pro-B cell ChIPseq. We would expect the reverse because accessibility cluster 19 is specific for progenitor cells. This deviation might reflect differences in cell state between the ChIPseq studies and Yoshida et al.

Figure 9B shows analogous results for EOMES ChIPseq peaks in the NK and CD8 thymocyte cell types. Our motif analysis showed enrichment of EOMES in co-clusters formed by locus clusters 1 and 8. Locus cluster 1 and 8 had enriched EOMES ChIPseq peak counts for both NK and CD8 thymocyte cell types and only the NK cell type, respectively. In-line with these results, early T cells - of which CD8 thymocytes are a member - and NK cells are both accessible in locus cluster 1 but only NK cells are accessible in locus cluster 8. Both NK and CD8 thymocytes had a statistically significant association between their accessibility co-clusters and the locus clusters at which EOMES peak counts were enriched (p-values 0.0003 and 0.001 respectively).

For both PAX5 and EOMES ChIPseq datasets, we also calculated the fraction of loci with peaks that were cell specific. Recall, cell specific loci were accessible in 2 or less of the Yoshida et al. cell types we considered. For PAX5 and EOMES, roughly 15% and 5% of loci with peaks were cell specific, respectively. In contrast, roughly 80% and 70% of loci with peaks fell within one of our 20 locus clusters, demonstrating that accessibility patterns across multiple cell types capture the dominant portion of TF binding, at least for EOMES and PAX5 and for the accessible loci we consider.

## 3 Materials and Methods

### 3.1 Construction of the *M* matrix

We downloaded the Yoshida fastq files from GEO accession GSE100738. We used the standard ENCODE ATACseq workflow to call peaks at different IDR as described in the text. We collected all peaks called by the ATACseq workflow across all cell types. Each peak was associated with a locus on the murine mm10 genome centered at the peak summit and extended 250 base pairs up and down stream. Then for each chromosome, moving 5′ to 3′, we sequentially evaluated the loci and formed a master list of loci. We did this in a greedy manner. If we encountered a locus that did not intersect with a locus already in our master list, we added the locus to the master list, otherwise we moved on to the next locus. Previous authors used a similar approach [20, 8, 43]. Given the master list of loci, we then formed *M*. A locus contained a peak for a given cell type if any of the cell types peaks intersected with the locus.

### 3.2 Locus Clustering

As an input graph to the Louvain algorithm, we let each row of *M* be a node and placed edges between rows (i.e. nodes) at a different FDR as described in the text. Each row consisted of 78 ones and zeros. Given two rows with *n* and *N* ones, respectively, we let *s* be the number of columns in which the two rows shared a 1. We assumed that *s* had a hypergeometric distribution (78 balls, *N* white balls and *n* draws), which corresponds to a null in which we permute the column of one of the rows. We then calculated the p-value cutoff that would lead to the specific FDR given the matrix *M*. Once the graph was constructed, we used the Python sklearn package implementation of the Louvain algorithm to perform the clustering.

### 3.3 Cell Type Clustering

On the differentiation tree, we defined a cell type cluster that respected the tree by a particular node *a*, which we called the cluster root node, and a subset of its children 𝒞_*a*_. The corresponding cluster contained all the nodes in 𝒞_*a*_ and a subset (possibly empty) of descendants of each of these nodes that formed a connected component of the tree. The root node *a* could be a member of the cluster, but could be kept out as well. This gave us a cluster that respected the tree, as described in the text. Partitioning the tree into *k* clusters is then equivalent to identifying *k* such *a*, 𝒞_*a*_ combinations. We took a greedy approach to finding the optimal *k* clusters that minimized 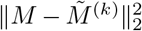. We started the algorithm by selected *k* root nodes randomly on the tree and for each root node *a*, including all its children in 𝒞_*a*_. Then at each iteration, we attempted to swap a root node with another node and attempted to swap the current subset of 𝒞_*a*_ with another subset. We repeated this iteration until no swaps improved the 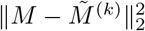. Since the problem is non-convex, we tried 20 different starting clusters for each *k*.

### 3.4 ANOVA Decomposition

For notational convenience, let *M* and 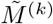 be the matrices *M* and 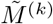 restricted to rows in a particular locus cluster. Then a standard ANOVA analysis decomposes the variation of *M* into a portion predicted by 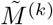 and a residual portion. The associated R-squared is given by,

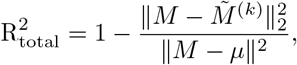

where *µ* is the mean of the entries of *M*. 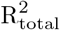 is exactly the standard R-squared of a linear predictor. The means of the columns of *M* are not affected by column clustering. With this in mind, we consider the prediction of the column means of *M*, which we write as *M*_*·,i*_, by 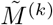. Then the portion of variation of *M*_*·,i*_ predicted by 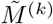 is given by the R-squared expression,

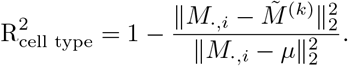

We calculate R-squared over the whole matrix *M* by averaging the R-squared values over the locus clusters.

### 3.5 TF Motif Analysis

Most of our TF analysis followed the workflow described in Schep et al. in [30]. We downloaded motif descriptions using the R package chromVARmotifs and then used the R package motifmatchR to call motifs on the DNA sequences spanned by our loci at a p-value of 5E-6. This gave us a binary matrix, *A*, analogous to *M* except that columns corresponded to motifs and a 0 and 1 value in an entry corresponded to the absence or presence of a motif at a locus, respectively.

To determine motif enrichment for a co-cluster, for each cell type in the co-cluster, we calculated the fraction of accessible loci that contained the motif and then averaged over all cell types in the cluster. This gave us a raw co-cluster score *r*. We computed an analogous raw null score *n* using all loci and cell types not in the co-cluster. Finally, we computed an enrichment score,

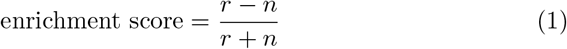

Schep et al. computed a similar score, except that they normalize *r* − *n* by a variance term. We found that the variance term was often small, leading to statistical instability, so instead we normalized by *r* + *n*. We then permuted columns to find a cutoff to our enrichment score that gave a 0.05 FDR. We called a motif as enriched for a co-cluster if its enrichment score exceeded the cutoff.

### 3.6 ChIPseq Workflow

We downloaded fastq files from Revilla-I-Doming et al., [28] with GEO accessions GSM932921, GSM932922 GSM932925, and GSM932926. We downloaded the fastq files from Wagner et al. [40] and Istaces et al. [17] with GEO accessions GSM3900380, GSM3900381 and GSM3559328, GSM3559327, respectively. We aligned the fastq to the murine mm10 genome using bowtie2 (-X1000 was the only non-standard flag) [19], filtered for poorly aligned reads and duplicates using samtools and Picard, and called peaks using MACS2 [45]. We called a loci from our ATACseq analysis as containing a ChIPseq peak if the ChIPseq peak summit was within the locus 500 base pair window.

## 4 Discussion

The recently published ATACseq dataset of Yoshida et al. provides a valuable resource through which to investigate patterns of chromatin accessibility across immune cell types. Here, we used this dataset to investigate the degree to which co-clustering of genomic loci and cell types can capture and describe patterns of chromatin accessibility. Some genomic loci were accessible in only 1 or 2 cell types, and we found that roughly half of accessible loci over immune cell types were of this type, in line with previous analyses of datasets encompassing non-hematopoietic cell types [38]. The other half of the accessible loci, which are accessible in multiple cell types, were the focus of our study. We found that essentially all of these loci, *>* 95%, can be grouped into 20 locus clusters. Within each locus clusters, the cell types showed roughly two states of accessibility reflecting a relatively high and low percentage of the loci that were accessible, respectively. For example, in locus cluster 1 which was composed of roughly 9000 loci, we found that in most T cell types, roughly 60% of the loci were accessible while in all other cell types roughly 0 − 10% were accessible. The dichotomy between cell types was more extreme in some locus clusters. For example, in locus cluster 8, we found all loci were accessible in NK cells but not in any other cell type. Ideally, in terms of cluster coherence, locus clustering might lead 100% or 0% of loci being accessible within a cell type. Certainly some of the locus cluster incoherence we see results from noise in the ATACseq workflow. But chromatin accessibility is not static, and some portion of the incoherence may reflect stochasticity in nucleosome positioning, binding of TF complexes, or other dynamic effects [41, 24]. It could also be that more locus clusters would result in greater coherence. Changing the graph we used as input to the Louvain clustering did not lead to more coherent locus clusters, but further work is needed to explore this issue.

Given the locus clusters, we found that a modest number of cell type clusters could capture a large fraction of the variation in accessibility associated with cell types. Using 12 cell type clusters, we were able to capture 80% of the cell type associated variation. Further, the cell type clusters we formed reflected coherent phenotypes as defined by the hematopoietic differentiation tree. When we formed cell type clusters using methods that were insensitive to the differentiation tree, the fraction of variance captured did not improve. Our cell type clustering extends the results of Lara-Astosia et al. [20] describing an association between accessibility and hematopoietic cell type.

Ultimately, we characterize chromatin accessibility to better understand cellular regulation. In particular, chromatin accessibility is strongly associated with TF binding [18]. Using both TF motif analysis and existing ChIPseq studies, we’ve shown that TF binding patterns associated with our co-clusters. Importantly, our results show that some TFs act across co-clusters. For example, we found that PAX5 motifs are enriched in two sets of loci. One set is accessible only in B cells while the other is accessible in both B cells and some myeloid cell types. Our ChIPseq analysis confirmed that PAX5 bound to both types of loci in B cells. Myeloid cell types do not express PAX5, at least at homeostasis, but the loci that are accessible in myeloid cell types and that are bound by PAX5 in B cells may regulate myeloid cells through other TFs or under non-homeostatic conditions. We found a different binding pattern for the transcription factor EOMES. In our ChIPseq analysis, we found that EOMES bound loci in NK and T cells, but that loci bound in NK cells were inaccessible in T cells and vice-versa. Our motif analysis suggests that many TF act across several co-clusters, in a manner similar to PAX5 and EOMES. These results suggest that analyzing patterns of chromatin accessibility through co-clustering or through a biclustering method may be essential in understanding the overlap and divergence of regulation in different cell types.

From a computational viewpoint, our work provided two insights. First, we found that co-clustering, rather than biclustering, provided a relatively stable and scalable means of analyzing ATACseq datasets across many cell types. We initially attempted a biclustering approach but found that solutions depended on starting conditions of the algorithm, that the algorithms did not scale well to the large number of accessible loci, and that interpretation was difficult. Second, we developed a novel graph based clustering algorithm to account for the hematopoietic differentiation tree. In this context, the novelty is the form that we assumed for the clusters. Initially, we formed clusters as connected components of the differentiation tree, but we found that the clusters created did not approximate the Yoshida et al. data well. Certain cell types have accessibility patterns that are different than the patterns of their parent cell type and connected components force the parent to be included with the children. Accounting for this effect vastly improved the fit of our clustering and points to the need for clustering approaches that account for the specifics of differentiation biology.

Our analysis involved several computational choices that may affect our results. We made binary calls of whether a locus was accessible or inaccessible. Using a continuous measure may better reflect chromatin accessibility biology and may affect our clustering results. From a computational perspective, we depended on the binary nature of the data to construct the input graph to the Louvain algorithm. We have also not explored an iterative co-clustering approach, e.g [6]. Our two-step clustering of loci followed by cell types makes our approach simple and scalable, but an iteration may lead to better results. Biologically, we are limited to the cell types given in the Yoshida et al dataset and our assumption of a particular form to the differentiation tree.

Overall, we have demonstrated a co-clustering approach that quantifies and delineates the association between chromatin accessibility and immune cell type. Our results provide a context in which to assess chromatin accessibility of other immune cell types. With the increased application of single cell ATACseq and the likely generation of even larger bulk ATACseq datasets, computational approaches to characterize chromatin accessibility patterns over an increasingly broad set of hematopoietic cell types will be needed.

## Acknowledgements

We thank the ImmGen consortium, without which this work would not have been possible.

## Notes

### Competing Interest Statement

The authors have declared no competing interest.

